## Supplementary material for "Ca^2+^ Oscillation in Vascular Smooth Muscle Cells Control Myogenic Spontaneous Vasomotion and Counteract Post-ischemic No-reflow": Supplmental Materials

**Inventory of Supplemental Materials**

Supplemental Figure Legends

Supplemental Movie Legends

Supplemental Figure S1-S13

#### Supplemental Figure Legends

**Fig. S1. Myogenic spontaneous vasomotion mainly governs cerebral arterioles but not venules in anesthetized mice.**

**a** and **b,** Maximal intensity projection (MIP) and still-frame images of the cerebral pial vessels, including the arteriole, venule, and penetrating arteriole (PA) in *SMACreER:Ai47* mice injected with RhoB intravenously under 2PLSM. **c,** Representative kymographs of spontaneous vasomotion in the bilateral arteriolar wall. **d,** Representative time-lapse bilateral arteriolar radius changes trace (dash and solid red line). **e,** Correlation coefficient analysis of bilateral arteriolar wall vasomotion, the magenta dash line indicates correlation coefficient is 0 (N = 7 mice, n = 30 vessels). **f,** Kymographs of spontaneous vasomotion in different vessels (corresponding resliced positions marked in **a** and **b**). The extravascular and luminal sides were distinguished by the RhoB blood signals (red) divided by the mural cell signals (green). **g,** Representative time-lapse radius changes trace of the arteriole (red), PA (orange), and venule (green). **h,** Fourier transform analysis of the rhythmic fluctuations in the pial arteriole, venule, and PA (N = 3 mice). **i,** Statistical analysis of accumulated power (AUC, area under the curve) of vasomotion within the frequency range of 0~0.3 Hz in the pial arteriole, venule, and PA (N = 3 mice). **j**-**l,** Vasomotion index analysis of arteriole(N = 3 mice, n = 15 or 29 vessels), venule (N = 3 mice, n = 6 or 17 vessels), and PA (N = 3 mice, n = 6 or 7 vessels), including frequency of rhythmic cycles (**j**), standard deviation (SD) of the peak intervals (**k**), and amplitude of changes in the vascular radius (**l**). Data are expressed as the mean ± SEM. Statistic for **i**, data were analyzed using one-way ANOVA, followed by Tukey's post hoc analysis. Statistics for **j**, **k**, and **l**, paired t-tests were used.

**Fig. S2. Arteriolar vasomotion index analysis between male and female mice. a-c,** Vasomotion index analysis between male (N = 6 mice, n = 38 vessels) and female (N = 5 mice, n = 33 vessels), including frequency of rhythmic cycles (**a**), standard deviation (SD) of the peak intervals (**b**), and amplitude of changes in the vascular radius (**c**). Data are expressed as the mean ± SEM. Unpaired t-tests were used for statistics.

**Fig. S3. Permanent unilateral CCA ligation (sham surgery) does not affect MCA spontaneous. a,** Experimental design of the sham surgery and timeline of 2PLSM imaging. **b,** Representative still frames of MCA before and after (sham 24h) sham surgery in *SMACreER:Ai47* mouse under 2PLSM, the vascular lumen was visualized through the intravenous injection of the red fluorescent dye rhodamine B-dextran. **c**-**e**, Representative kymographs (corresponding resliced positions marked in **b** of blood vessels and vasomotion index analyses before and after (sham 24h) sham surgery, including arteriole (**c**), PA (**d**) and venule (**e**) (N = 2 mice, n = 6~29 vessels). **f,** LSCI images of the mouse whole brain indicate the time course changes in the CBV after sham surgery. **g,** Statistical analysis of the relative CBV changes at different time points after sham surgery (N = 5 mice). All measurements were normalized to the basal CBV before surgery. Data are expressed as the mean ± SEM. Statistics for **c**, **d** and **e**, unpaired t-tests were used.

**Fig. S4. MCAO model induces mild capillary obstruction in the time point of 2h occlusion and 22h reperfusion. a,** Representative frame-scan images of RhoB-labeled capillary blood flow before and after ischemic stroke. The turquoise dots represent the spatial position of RBCs. **b,** Capillary stall rate analysis during MCAO-induced ischemic stroke, which was calculated as the percentage of stalled capillaries (no blood cell flowing longer than 10.84 seconds/10 frames) among all capillaries under one imaged view (0.259 $mm^2$) (N = 8 mice).

**Fig S5. MCAO does not affect brain vasculature density in the time point of 2h occlusion and 22h reperfusion. a,** Representative images of whole brain vessels in brain slices after 2h occlusion and 22h reperfusion. Brain vessels (including arterioles, capillaries, and venules) were viewed using *Cdh5CreER:Ai47* mice, which were used to label endothelial cells with enhanced green fluorescent protein (EGFP). Hoechst 33342 (magenta) was used to label nuclei, which exhibited condensed morphology in the infarct core of ipsilateral hemisphere (white arrow). **b,** Statistical analysis of the capillary density between the contralateral and ipsilateral sides after 2h occlusion and 22h reperfusion in the MCAO model (N = 3 mice, n= 18 views). **c,** Representative image of arteriolar vasculature on the surface of the entire brain after 2h occlusion and 22h reperfusion. The brain arterioles were reported using *SMACreER:Ai47*. **d,** Statistical analysis of the pial arteriole density between the contralateral and ipsilateral sides after 2h occlusion and 22h reperfusion in the MCAO model (N = 5 mice). Data are expressed as the mean ± SEM. Data were analyzed using unpaired t tests in **b** and paired t-tests in **d**.

**Fig S6. MCAO does not affect SMC integrity of adult mice in the time point of 2h occlusion and 22h reperfusion in vivo. a,** Representative MIP images of MCA in the time point before and after (Occ.2hRep.22h) occlusion in live mice under 2PLSM. Using the *SMACreER:Ai47* mice, SMCs were reported by the green fluorescent EGFP after tamoxifen administration. The labeling rate of SMCs (SMCs were not fully labeled as discontinued fluorescent SMCs along MCA in the present data) could be modified by increasing the tamoxifen dosage or the induction time. As the EGFP is expressed in the cytosol of SMCs, the retaining of the signal could manifest the membrane integrity and the live state of SMCs. The labeled SMCs were identical before and after ischemic stroke. **b,** Statistical analysis of the SMC density along the arterioles showed no significant difference between the two time points (N = 5 mice, n = 5 arterioles). Data are expressed as Mean ± SEM. Data were analyzed using paired t-test.

**Fig S7. Ischemic stroke evokes transient ΔΨm loss in SMCs. a,** Experimental design of the ex vivo SMC ΔΨm detection under 2PLSM. **b,** Representative images of TMRM labeled MCA of the contralateral and ipsilateral hemisphere after 2h occlusion and 1h reperfusion ex vivo. The dashed lines circle the MCA area. Green arrowheads represent blood flow direction. **c,** Relative TMRM intensity analyses between the contralateral and ipsilateral MCA after 2h occlusion and 1h reperfusion ex vivo (N = 6 mice, n= 6 views). **d,** Representative images of TMRM labeled MCA of the contralateral and ipsilateral hemisphere after 2h occlusion and 22h reperfusion ex vivo. The dashed lines circle the MCA area. Green arrowheads represent blood flow direction. **e,** Relative TMRM intensity analyses between the contralateral and ipsilateral MCA after 2h occlusion and 22h reperfusion ex vivo (N = 4 mice, n= 8 views). **f,** Time course analysis of the relative ΔΨm changes in SMCs of the MCA during ischemic stroke, combined with the data in vivo and ex vivo. Data are expressed as Mean ± SEM. Data were analyzed using unpaired t-test.

**Fig S8. IP3R inhibition with 2-APB suppress calcium oscillation in primary SMCs.** **a,** Experimental timeline of live imaging before and after 2-APB (10μM) in Fluo-4 loaded primary SMCs. **b,** Still-frame images and kymographs of primary-culture SMCs (red, Tdt+) before and after 2-APB (10μM) treatment. The calcium signal was indicated by Fluo-4. The magenta dots in kymographs indicate the calcium oscillation peaks in the ROI area (yellow line). **c,** Representative time-lapse fluorescent Fluo-4 signal changes trace in the cytoplasm of primary-culture SMCs before and after 2-APB (10μM) treatment. **d,** Statistical analysis of the calcium oscillation frequency and amplitude before and after 2-APB (10μM) treatment in primary-culture SMCs (N = 3 assays, n = 40 cells). Data are expressed as the mean ± SEM. Statistics were analyzed using paired t-tests.

**Fig S9. The establishment and targeting strategy of RCL-*ME-Linker* mice**. **a,** RCL (ROSA26/CAG promoter/*LoxP-STOP-LoxP*)-*ME-Linker* mice were generated by inserting the targeting elements into the ROSA26 locus in C57BL/6N lines. The targeting elements contain a CAG promoter, a floxed terminator, and the core targeting gene *Mito-ER-linker-P2A-GCaMP6s*. The *P2A*-linked *GCaMP6s* gene was designed to report cytosolic calcium signals in ME-Linker-overexpressing cells. **b,** In the presence of Cre enzyme, the *STOP* gene was excised, thereby allowing the CAG promoter to drive *ME-Linker* and *GCaMP6s* expression. *ME-Linker* mice were crossed with Cre mice (specific cell or tissue Cre enzyme expression) to obtain double-positive mice; in this way, ME-Linker and the calcium indicator GCaMP6s could be overexpressed in the targeted cells in vivo.

**Fig S10. ME-Linker overexpression in SMCs does not affect cardic functions, blood pressure, and body temperature. a,** Representative echocardiographic images in B-Mode and M-Mode of the left ventricular (LV) in control littermates and *SMAcreER:ME-Linker* mice. **b-m,** Statistical analysis of heart rate (**b**), LV mass (**c**), LV ejection fraction (**d**), LV fractional shortening (**e**), cardiac output (**f**), stroke volume (**g**), and ventricular end-systolic (;s) or end-diastolic (;d) left ventricular interior diameter (LVID) (**h** and **i**), left ventricular posterior wall (LVPW) (**j** and **k**), and left ventricular anterior wall (LVAW) (**l** and **m**) in control littermates and *SMAcreER:ME-Linker* mice. **n,** Representative echocardiographic images in PW Doppler-Mode and PW tissue-Mode in detecting mitral valve (MV) blood velocity and MV velocity. **o,** Statistical analysis of MV function by calculating E/E' values in control littermates and *SMAcreER:ME-Linker* mice. **p,** Representative echocardiographic images in PW Doppler-Mode of aortic arch (Ao Arch), aortic valve (AV) and pulmmonary valve (PV). **q-s,** Statistical analysis of Ao Arch (**q**), AV(**r**) and PV(**s**) peak velocity in control littermates and *SMAcreER:ME-Linker* mice. N = 4 mice for each genotypes in echocardiographic assays. **t,** Analysis of systolic blood pressure in control littermates and *SMAcreER:ME-Linker* mice before and after 1 week of tamoxifen treatment (N = 3 or 5 mice). **u,** Analysis of body temperature in control littermates and *SMAcreER:ME-Linker* mice before and after 1 week of tamoxifen treatment (N = 5 or 7 mice). **v-x,** Vasomotion index analysis between control littermates (N = 6 mice, n = 38 vessels) and *SMAcreER:ME-Linker* mice (N = 5 mice, n = 33 vessels), including frequency of rhythmic cycles (**v**), standard deviation (SD) of the peak intervals (**w**), and amplitude of changes in the vascular radius (**x**). Data are expressed as the mean ± SEM. All data were analyzed using unpaired t-tests.

**Fig S11. ME-Linker overexpression in SMCs does not affect neuronal survival threshold to ischemic injury. a,** Experimental timeline of the in vitro oxygen-glucose deprivation (OGD) assay of acute brain slices. The acute brain slices from control (*ME-Linker*) and *SMACreER:ME-Linker* mice were prepared quickly in half an hour. Afterwards, 1h of control or OGD condition was exposed to the corresponding slices, and the threshold to ischemic injury was detected through PI staining after PFA fixing. **b,** Representative images of PI (red) and Hoechst33342 (turiquoise) staining results in control and OGD brain slices of control (*ME-Linker*) and *SMACreER:ME-Linker* mice. **c,** Statistical analysis of the brain cell death percentage (proportation of PI positive nueleus in Hoechst33342 positive nucleus) in control and OGD brain slices of control (*ME-Linker*) and *SMACreER:ME-Linker* mice (N = 4 slices). **d,** Representative images of PI (red) and NeuroTrace Fluorescent Nissl (turiquoise) staining results in control and OGD brain slices of control (*ME-Linker*) and *SMACreER:ME-Linker* mice. **e,** Statistical analysis of the neuron death percentage (proportation of PI positive nueleus in Nissl positive nucleus) in control and OGD brain slices of control (*ME-Linker*) and *SMACreER:ME-Linker* mice (N = 4 slices). **f,** Representative images of Hoechst33342 (green) and NeuroTrace Fluorescent Nissl (magenta) staining results in control and OGD brain slices of control (*ME-Linker*) and *SMACreER:ME-Linker* mice. **g,** Statistical analysis of the neuron percentage (proportation of Nissl positive nueleus in Hoechst33342 positive nucleus) in control and OGD brain slices of control (*ME-Linker*) and *SMACreER:ME-Linke*r mice (N

= 4 slices). Data are expressed as the mean ± SEM. Statistics were analyzed using unpaired t-tests.

**Fig S12. Neutrophil depletion do not helps in restoring no-reflow in *SMACreER:ME-Linker* mice. a,** Scheme of timepoints for MCAO surgery, CBF detection using LSCI, NSS detection, and histological assessment in neutrophil depletion assay. The antibody or rt-PA administration and hematology analysis were performed in the corresponding timepoint as appropriate. **b,** Scatter plots of the leukocyte classification in the peripherial whole blood using ProCyte Dx hematology analyzer between control (*ME-Linker*) and *SMACreER:ME-Linker* mice. **c,** Statistical analysis of the proporation of neutrophil (NEUT), lymphocyte (LYMPH), monocyte (MONO), eosinophil (EO), and basophil (BASO) in total number of leukocytes count, before Ly6G antibody and isotype antibody administration. **d,** The absolute number (K/μl) of all kinds of leukocyte in the peripherial whole blood in different groups, before Ly6G antibody and isotype antibody administration. **e,** Scatter plots of the leukocyte classification in the peripherial whole blood using ProCyte Dx hematology analyzer at the time point of reperfusion 20h after ischemic stroke, in anti-Ly6G antibody and isotype antibody administrated mice. **f,** Statistical analysis of the proporation of different leukocytes in total number of leukocytes count, after Ly6G antibody and isotype antibody administration. **g,** The absolute number (K/μl) of all kinds of leukocyte in the peripherial whole blood in differents groups, after Ly6G antibody and isotype antibody administration. **h,** LSCI images of the mouse whole brain indicate the time course changes in the cerebral CBF before and after MCAO surgery in different groups. **i,** Statistical analysis of the relative cerebral CBF changes at different time points before and after MCAO surgery in different groups (N = 7 or 8 mice for each groups). All measurements were normalized to the basal cerebral CBF before surgery. Data are expressed as the mean ± SEM. Statistics for **g**, one-way ANOVA, followed by Tukey's post hoc analysis were used. Statistics for **d** and **i**, by using unpaired t test, p values (*) were evaluated between control (*ME-Linker*) and *SMACreER:ME-Linker* mice at the timepoint of Occ.2hRep.22h.

**Fig S13. ME-Linker overexpression in SMCs attenuate behavioral and histological injuuries after ischemic stroke. a,** Scheme of of timepoints for MCAO surgery, NSS (neurological severity score) detection, and histological assessment in the long-term (2 weeks) of ischemic-induced neuronal injury evaluation assay. **b,** Statistical analysis of the NSS along two weeks after ischemic stroke in control (*ME-Linker*) and *SMACreER:ME-Linker* mice (N = 3 mice for each group). **c,** Heat map of inspection items included in the NSS examination along days after ischemic stroke, as shown with control (*ME-Linker*) mice in blue, and *SMACreER:ME-Linker* mice in red. **d,** Representative images of the mouse brains after ischemic stroke in control (*ME-Linker*) and *SMACreER:ME-Linker* mice in the timepoint of two weeks after ischemic stroke. **e,** Statistical analysis of the cortical hemisphere atrophy rate after two weeks ischemic stroke in control (*ME-Linker*) and *SMACreER:ME-Linker* mice (N = 3 mice). **f,** (Left) Representative images of Map2 staining results in two weeks post-ischemic brain slices, the bountry of the ischemic area (magenta dotted line) was distinguished by the positive signal of Map2 labeled neuronal processes. (Right) Representative serial section of Map2 staining slices in control (*ME-Linker*) and *SMACreER:ME-Linker* mice after two weeks ischemic stroke. **g,** Statistical analysis of the neuronal injury (Map2 loss) volume after two weeks ischemic stroke in control (*ME-Linker*) and *SMACreER:ME-Linker* mice (N = 3 mice). **h,** (Left) Representative images of NeuN staining results in two weeks post-ischemic brain slices, the bountry of the ischemic area (magenta dotted line) was distinguished by the positive signal of NeuN labeled neuronal somas. (Right) Representative serial section of NeuN staining slices in control (*ME-Linker*) and *SMACreER:ME-Linker* mice after two weeks ischemic stroke. **i,** Statistical analysis of the neuronal injury (NeuN loss) volume after two weeks ischemic stroke in control (*ME-Linker*) and *SMACreER:ME-Linker* mice (N = 3 mice). **j,** (Left) Representative images of NeuroTrace Fluorescent Nissl staining results in two weeks post-ischemic brain slices, the bountry of the ischemic area (magenta dotted line) was distinguished by the breakage signal of Nissl substance around neuronl somas. (Right) Representative serial section of NeuroTrace Fluorescent Nissl staining slices in control (*ME-Linker*) and *SMACreER:ME-Linker* mice after two weeks ischemic stroke. **k,** Statistical analysis of the neuronal injury (Nissl breakage) volume after two weeks ischemic stroke in control (*ME-Linker*) and *SMACreER:ME-Linker* mice (N = 3 mice). Data are expressed as the mean ± SEM. Statistics were analyzed using unpaired t-tests.

#### Supplemental Movie Legends

**Movie 1.** 2PLSM time-lapse imaging of the cerebral pial vessels, including the arteriole, venule, and penetrating arteriole (PA) in *SMACreER:Ai47* mice injected with RhoB intravenously under 2PLSM.

**Moive 2.** Time-lapse imaging of MCA before and after (occ.2hrep.22h) ischemic stroke in *SMACreER:Ai47* mouse under 2PLSM. Coupled with the real-time radius changes trace of arteriole before (dash line) and after (solid line) (occ.2hrep.22h) ischemic stroke.

**Moive 3.** Time-lapse imaging of MCA before and after (occ.2hrep.22h) ischemic stroke in SMACreER:Ai96 mouse under 2PLSM. Coupled with the real-time calcium oscillation changes trace of SMC before (dash line) and after (solid line) (occ.2hrep.22h) ischemic stroke.

**Moive 4.** Time-lapse imaging of primary-culture SMCs before and after CCCP treatment in the control virus and *ME-Linker* virus groups. The calcium signal was indicated by calcium indicator YTnC2-5.

**Moive 5.** Time-lapse imaging of calcium oscillation changes in SMC before and after (occ.2hrep.22h) ischemic stroke in *SMACreER:ME-Linker* mouse under 2PLSM. Coupled with the real-time calcium oscillation changes trace of SMC before (dash line) and after (solid line) (occ.2hrep.22h) ischemic stroke.

**Moive 6.** Time-lapse imaging of MCA before and after (occ.2hrep.22h) ischemic stroke in *SMAcreER:ME-Linker* mice mouse under 2PLSM. Coupled with the real-time radius changes trace before (dash line) and after (solid line) (occ.2hrep.22h) ischemic stroke.

Supplementary Figure 1

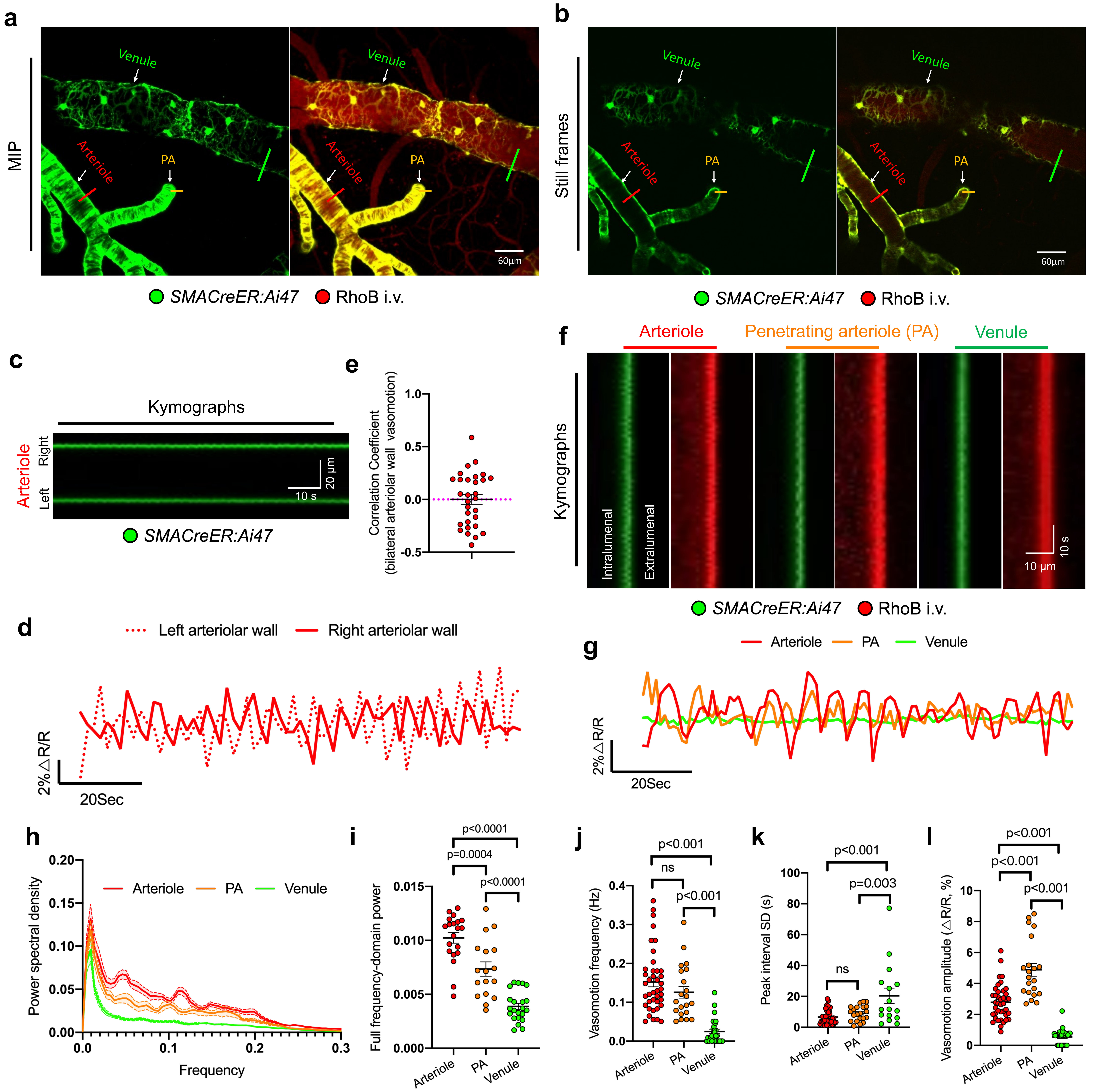

Supplementary Figure 2

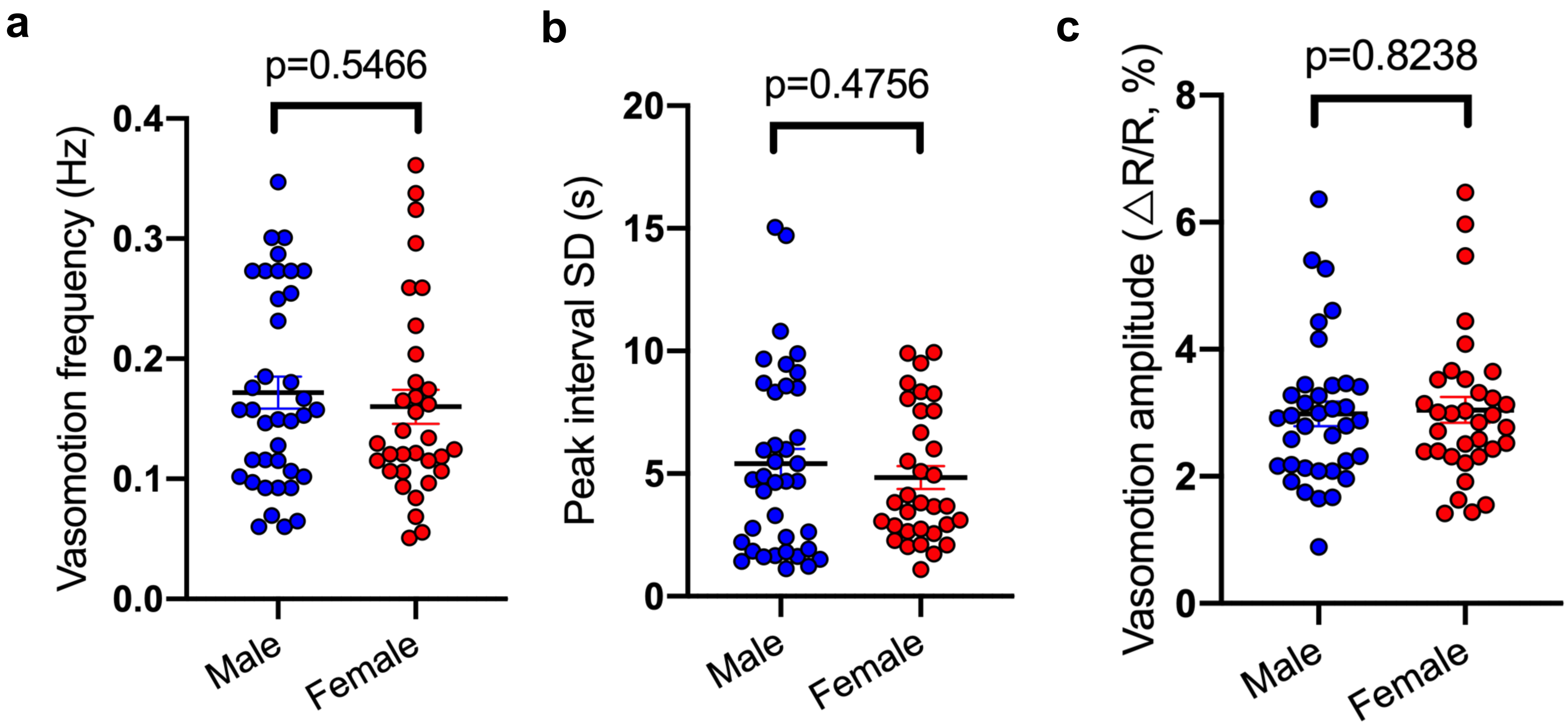

Supplementary Figure 3

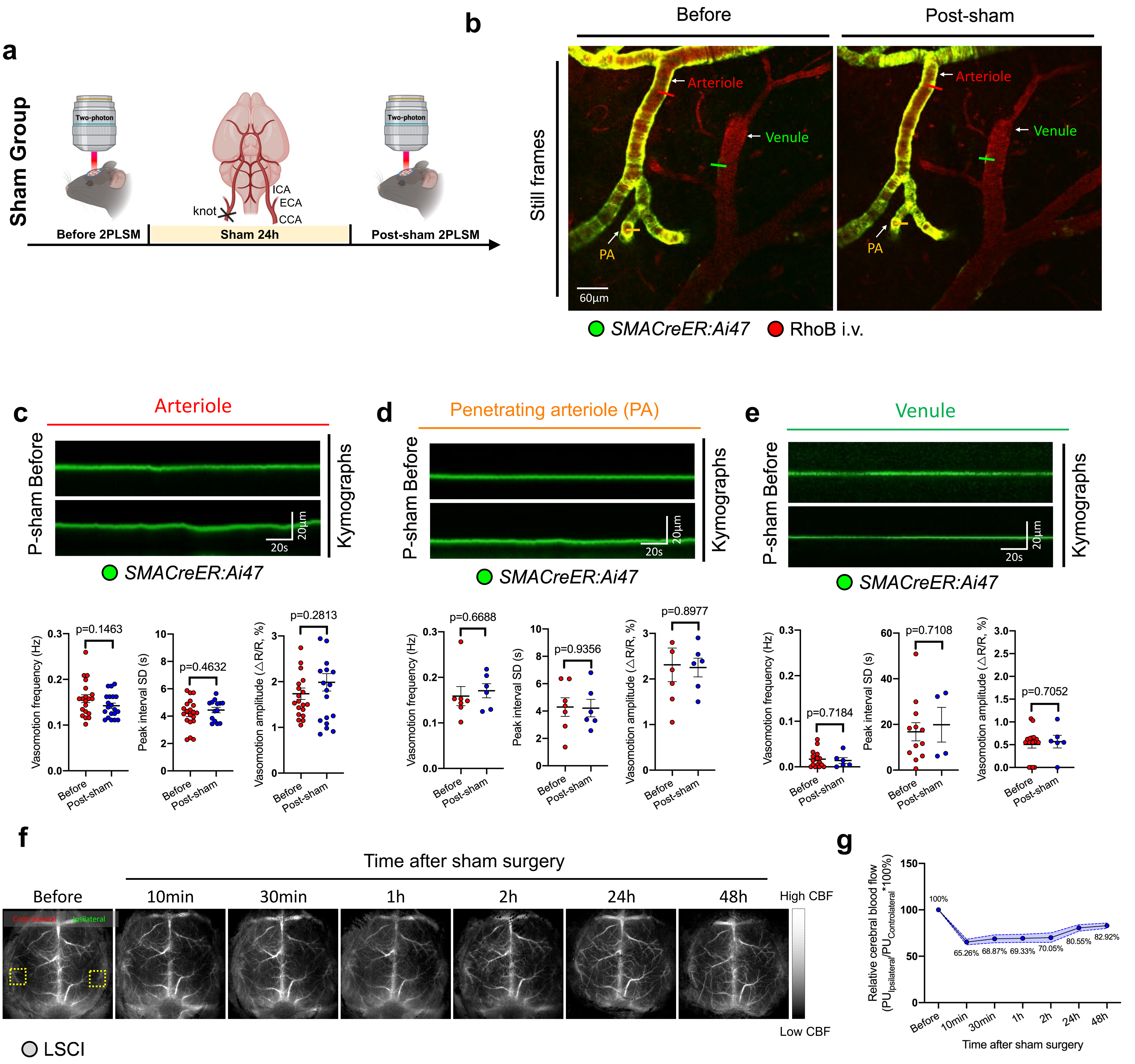

Supplementary Figure 4

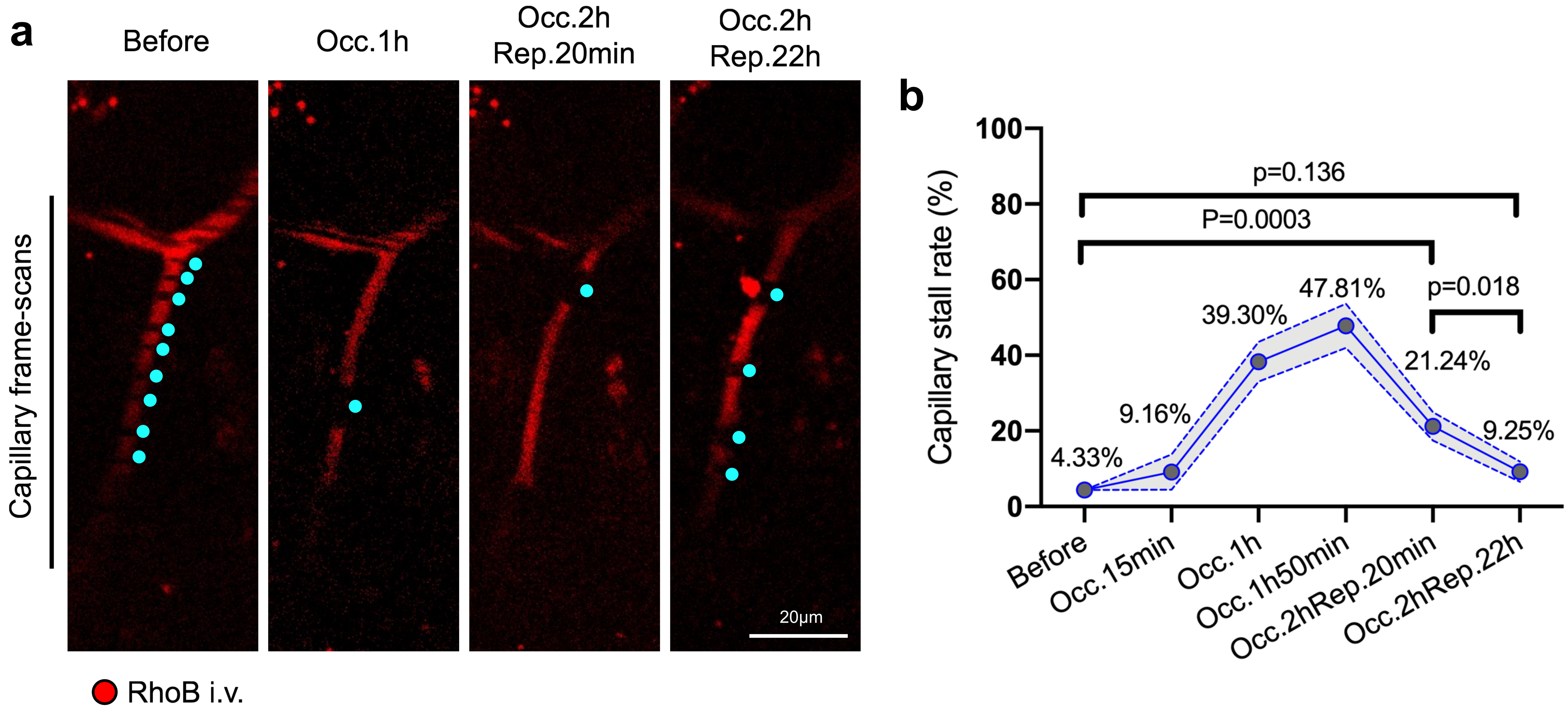

### Supplementary Figure 5

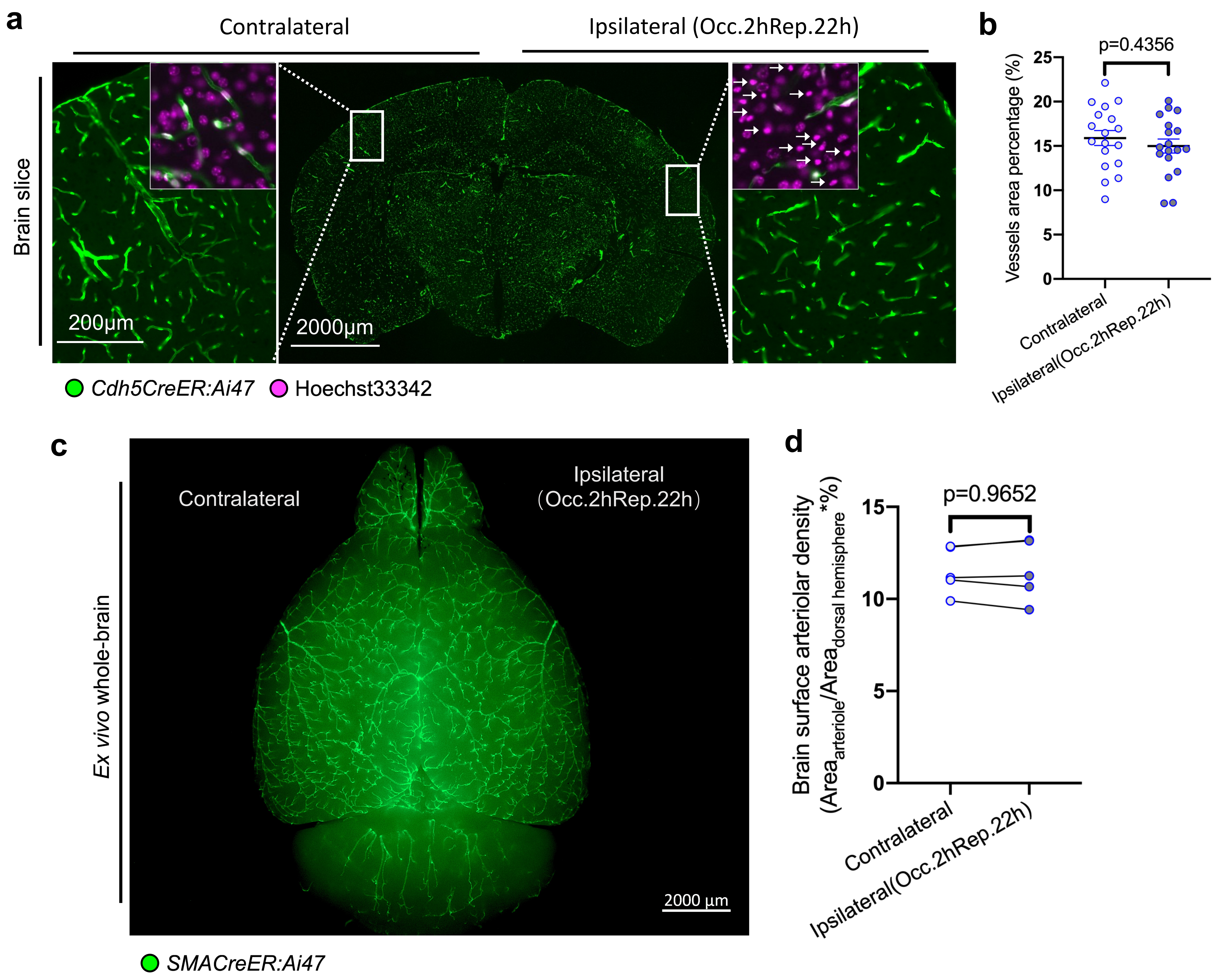

Supplementary Figure 6

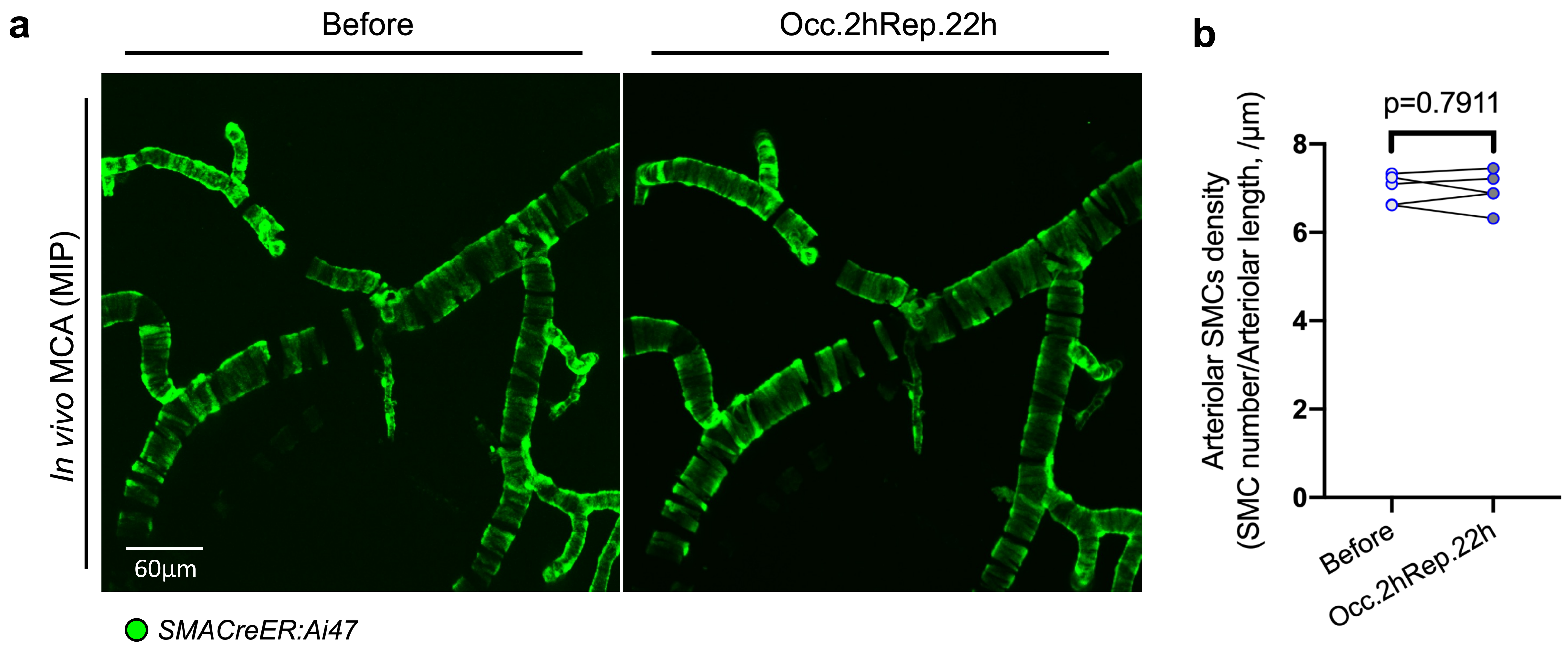

Supplementary Figure 7

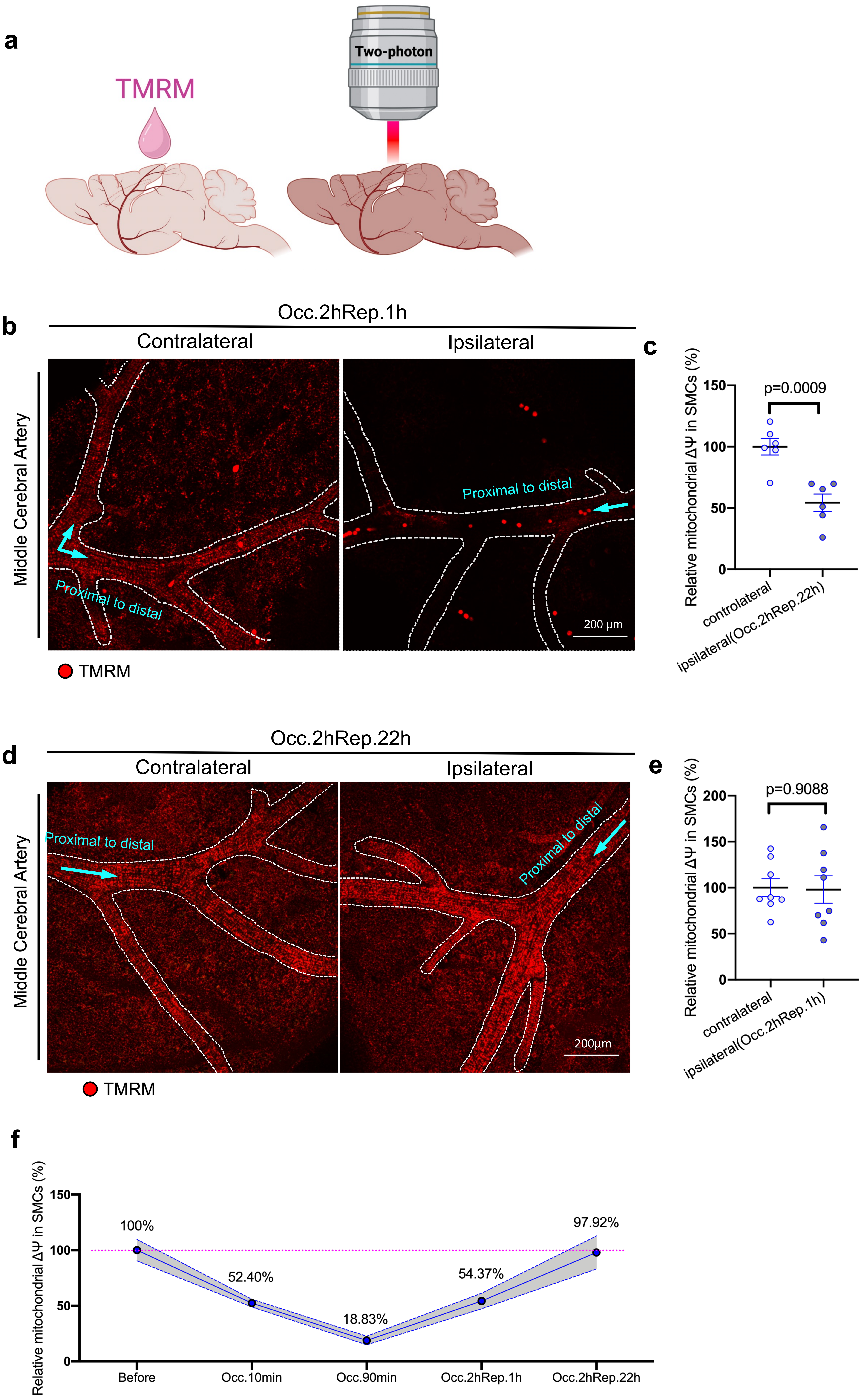

Supplementary Figure 8

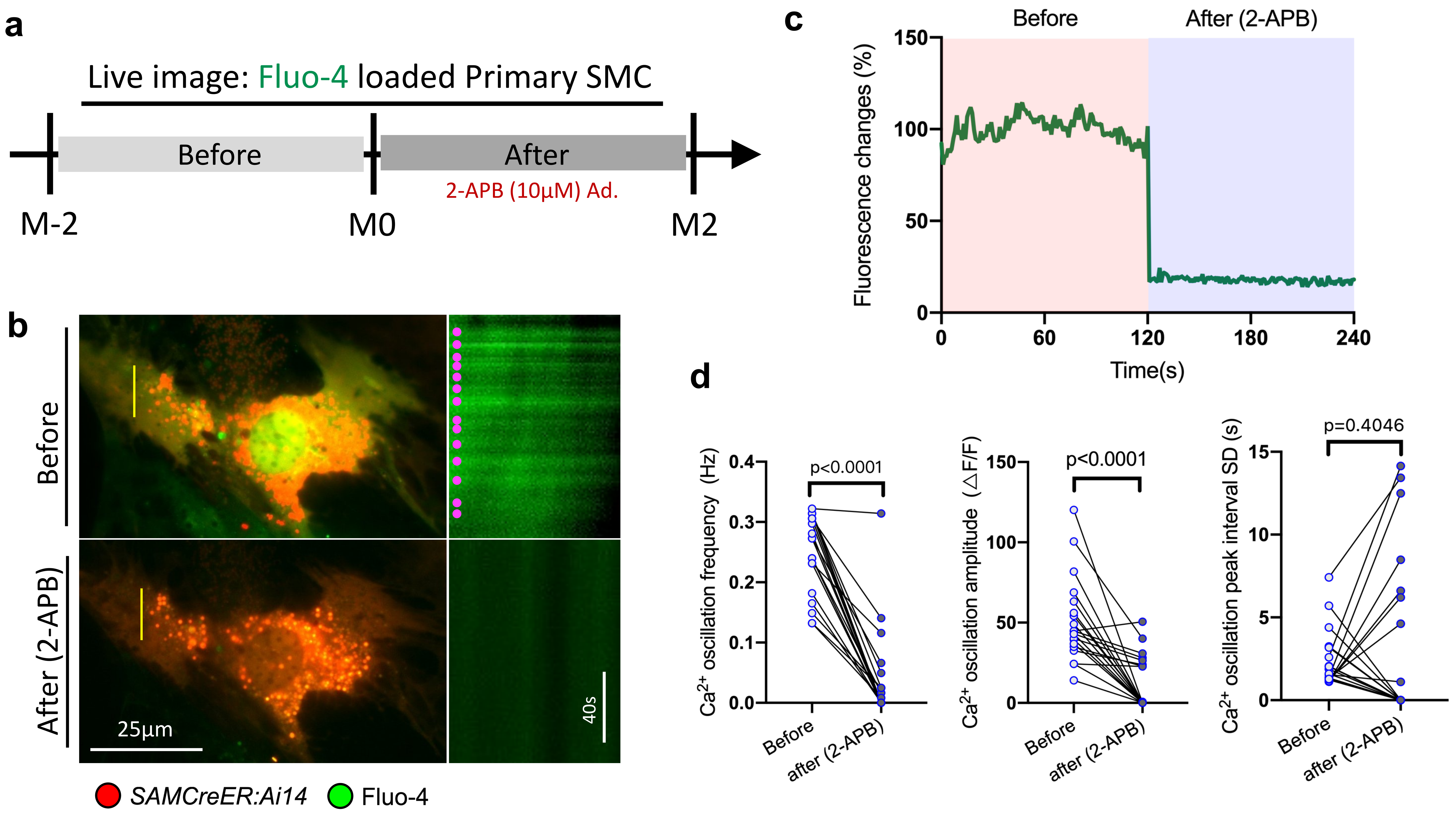

### Supplementary Figure 9

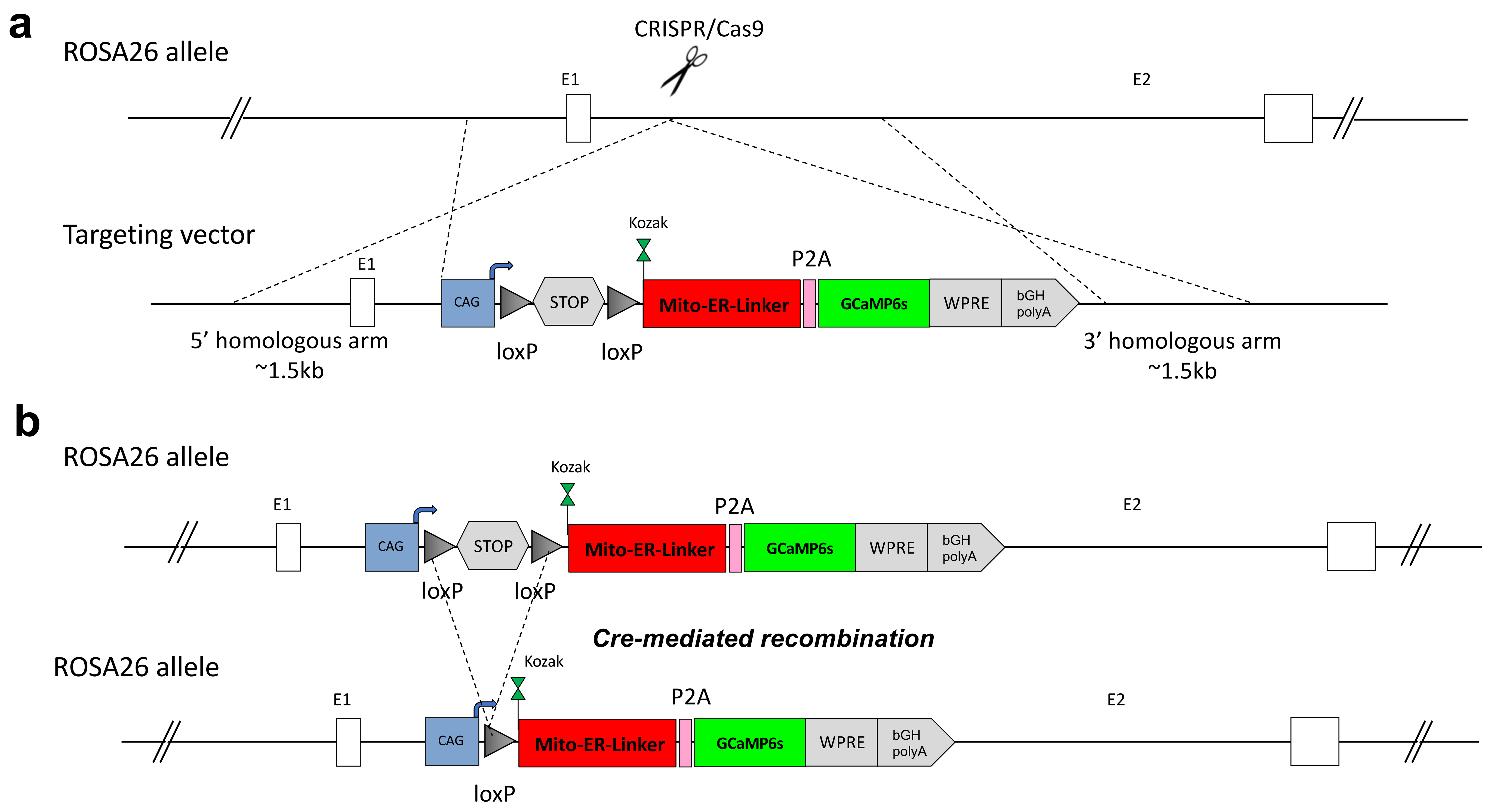

### Supplementary Figure 10

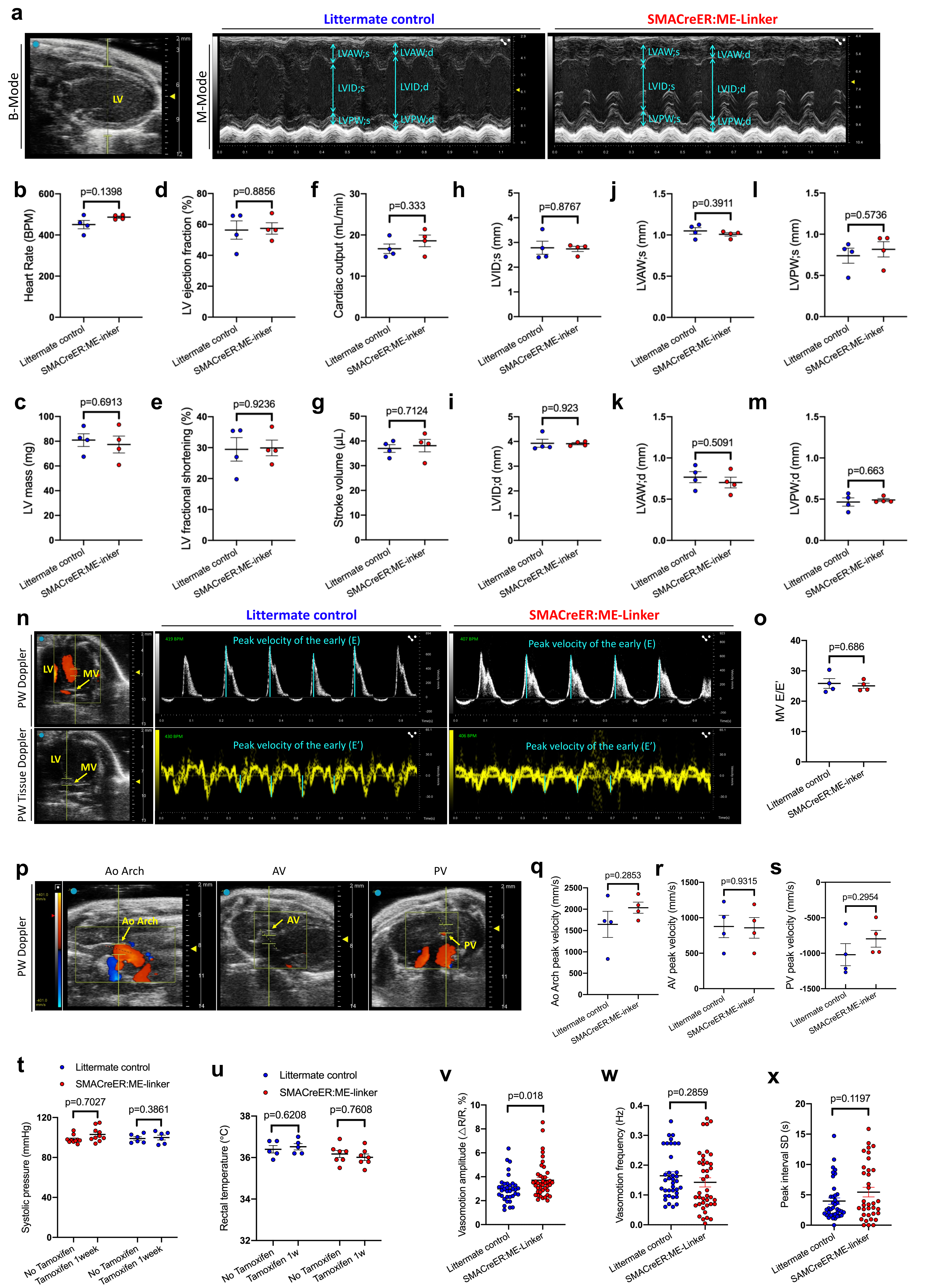

### Supplementary Figure 11

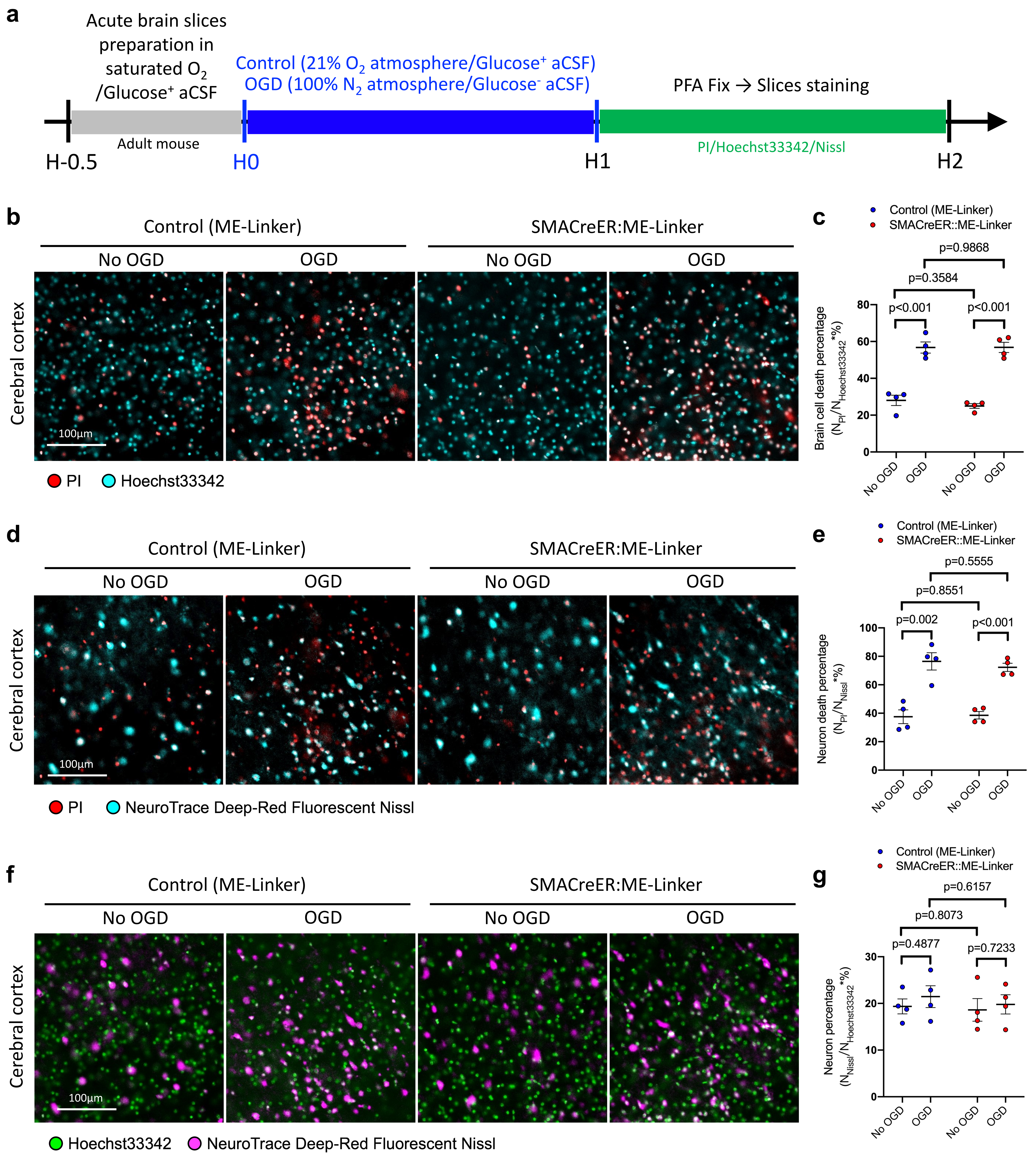

Supplementary Figure 12

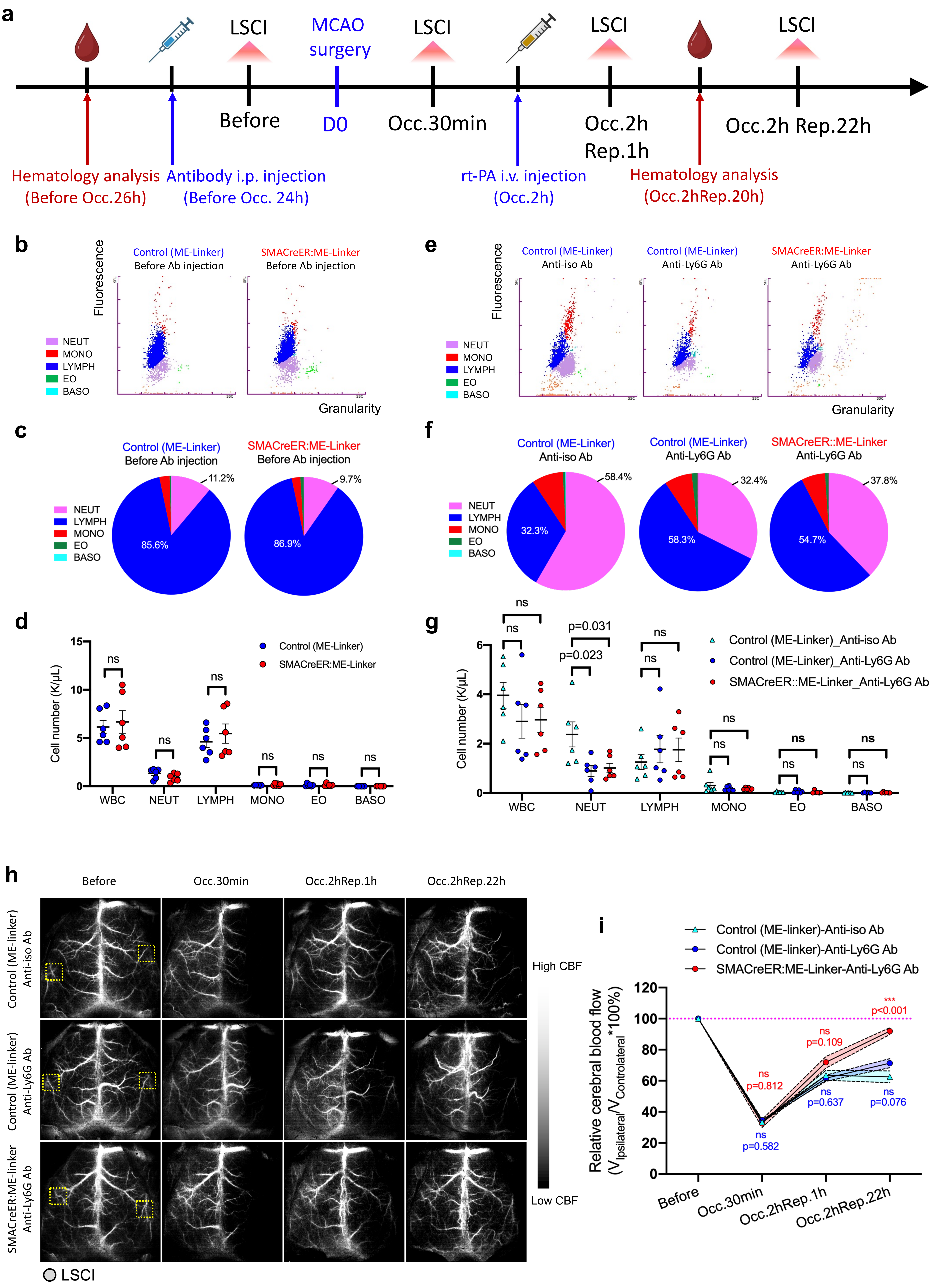

Supplementary Figure 13

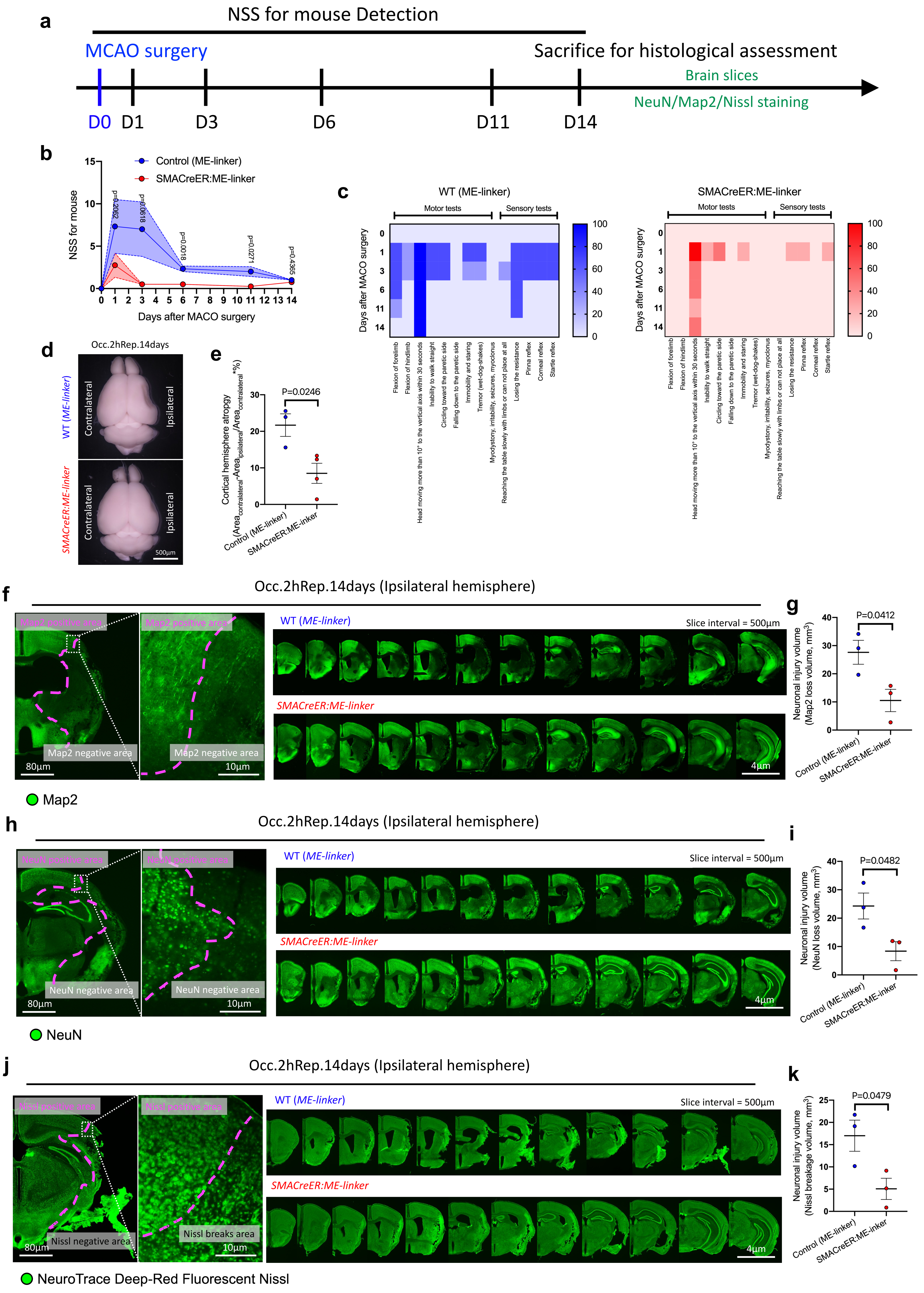
